# Mobile ear-EEG to study auditory attention in everyday life

**DOI:** 10.1101/2020.09.09.287490

**Authors:** Daniel Hölle, Joost Meekes, Martin G. Bleichner

## Abstract

Most research investigating auditory perception is conducted in controlled laboratory settings, potentially restricting its generalizability to the complex acoustic environment outside the lab. The present study, in contrast, investigated auditory attention with long-term recordings (*>*6 h) beyond the lab using a fully mobile, smartphone-based ear-centered electroencephalography (EEG) setup with minimal restrictions for participants. Twelve participants completed iterations of two variants of an oddball task where they had to react to target tones and to ignore standard tones. A rapid variant of the task (tones every 2 seconds, 5 minutes total time) was performed seated and with full focus in the morning, around noon and in the afternoon under controlled conditions. A sporadic variant (tones every minute, 160 minutes total time) was performed once in the morning and once in the afternoon while participants followed their normal office day routine. EEG data, behavioural data, and movement data (with a gyroscope) were recorded and analyzed. The expected increased amplitude of the P3 component in response to the target tone was observed for both the rapid and the sporadic oddball. Miss rates were lower and reaction times were faster in the rapid oddball compared to the sporadic one. The movement data indicated that participants spent most of their office day at relative rest. Overall, this study demonstrated that it is feasible to study auditory perception in everyday life with long-term ear-EEG.

---

Most research on the neural processes underlying auditory perception is conducted in the lab. The acoustic environment we are exposed to in daily life, however, is more complex than the controlled stimuli that are used in the lab. We are surrounded by several sound sources simultaneously, we move relative to these sound sources, we hear sounds while we are engaged in other tasks, and we shift our attention between different sound sources. Some sounds, such as a cell phone call, require our immediate reaction; other sounds we try to ignore. Complex task demands can cause us to miss a loud alarm. A pilot, for example, can miss an auditory alarm when the task demands of aviating are high (Dehais, Roy, & Scannella, 2019). In other cases, the persistent soft trickle of a dripping faucet can distract us profoundly. Clearly then our auditory perception is context dependent. A comprehensive understanding of auditory perception therefore requires study of the underlying neural processes in everyday life.

With new mobile electroencephalography (EEG) hardware using either electrodes on the scalp (Gramann, Ferris, Gwin, & Makeig, 2014; Gramann et al., 2011), around the ear (Bleichner & Debener, 2017; Debener, Emkes, De Vos, & Bleichner, 2015) or even inside the ear (Kidmose, Looney, Ungstrup, Rank, & Mandic, 2013; Looney et al., 2011), it is possible to leave the lab and to study neural processes beyond the lab while people are walking (De Sanctis, Butler, Malcolm, & Foxe, 2014; Debener, Minow, Emkes, Gandras, & de Vos, 2012), cycling (Scanlon, Townsend, Cormier, Kuziek, & Mathewson, 2019), working (Wascher et al., 2016), or aviating (Dehais, Duprès, et al., 2019). We can now study brain dynamics during the course of a day and in relation to different contexts using long-term mobile EEG recordings. In fact, past research has shown that cognitive performance and brain responses to auditory events fluctuate during the course of the day (Aseem & Hussain, 2019; Basinou, sub Park, Cederroth, & Canlon, 2017). These natural fluctuations are usually not considered in classical lab experiments, but could be captured with long-term EEG recordings. Fatigue effects during working or driving could also be investigated (Wascher et al., 2019, 2016).

However, there are technical and methodological challenges to overcome. When leaving the lab, increased movement artifacts may interfere with interpretation of the data (Ladouce, Donaldson, Dudchenko, & Ietswaart, 2017). We have to strike a balance between a natural experience for the participant to study real-life auditory perception and sufficient experimental control to draw meaningful conclusions. Furthermore, EEG analysis requires accurate identification of well-defined events. Finally, we should avoid to produce detached knowledge with this new technology, but should build on the knowledge that was obtained in lab-based research. This way we can deduce expectations from prior research and evaluate how they generalize to everyday contexts.

The objective of the present study is to demonstrate the feasibility of long-term EEG measurements of auditory attention in office workers during their workday. For this we used an adaptation of the well-studied oddball task. In this paradigm, series of frequent standard stimuli contain rare target stimuli (oddballs) which elicit an event-related potential (ERP) component, the P3, with a latency of 250-500 ms (Polich, 2007). The P3 is considered an index of auditory attention and can be captured in a lab-based oddball task using ear-EEG (Debener et al., 2015), but also in more natural settings, for example, while walking (Debener et al., 2012), cycling (Scanlon et al., 2019), or aviating (Dehais, Duprès, et al., 2019). We adapted the oddball task in such a way that it could be done in parallel to normal office work. To monitor the participants brain response to the task events during a full workday (>6 h) we used a mobile ear-EEG (cEEGrid, see Bleichner & Debener, 2017) setup.

In the current study, participants performed two variants of the oddball task where they were presented with a sequence of tones. Some of these tones served as target and required a response while the other tones did not. The rapid task was the classical oddball task with a quick succession of tones (Polich, 2007). Participants had to continuously concentrate on the sound sequence and react to target tones as quick as possible. The sporadic task, a version of the rapid task with longer inter-stimulus intervals, was intended for the office recording and to be performed in parallel to normal office work. While the rapid task lasted about 5 minutes, in the sporadic task the same number of stimuli was distributed over 160 minutes. We hypothesized that the target tone would elicit a P3 response in both oddball variants (Polich, 2007).

As mentioned before, extensive movements are detrimental to EEG signal quality and may impede valid conclusions. While numerous algorithms exist to deal with movement artifacts (e.g., Blum, Jacobsen, Bleichner, & Debener, 2019; Oliveira, Schlink, Hairston, König, & Ferris, 2016,), the best EEG results can be expected when participants barely move. For this reason, participants are asked to sit as still as possible in a classical EEG lab study. When moving beyond the lab, we can expect to see more artifacts due to participant movement. However, beyond the lab EEG acquisition does not necessarily imply EEG acquisition during movement. For example, you might realize now, that you have not moved much while reading this text. We expected that during an office day people spend considerable time with only minor movement. To investigate this intuition, we recorded the gross body movement of participants to identify movement-free periods throughout the experiments. Quantifying the portion of data that is (not) artifact-contaminated can help to predict data quality in future research.

Behaviorally, we predicted faster reaction times in rapid than in sporadic oddball blocks. Similarly, we predicted fewer misses in rapid compared to sporadic oddball blocks. These predictions were based on the assumption that participants would be more distracted by other tasks while following their daily work routine during sporadic oddball blocks than during the rapid oddball blocks, where they focused on the oddball task.

## Methods

### Participants

Twelve healthy volunteers (7 female, 5 male; 2 left-handed, 9 right-handed, 1 ambidextrous) with self-reported normal hearing were recruited by word-of-mouth advertising. Participants were either undergraduate students or staff members of the Oldenburg Department of Psychology and had previous experience with EEG. Their age ranged from 24 to 39 (*M* = 27.83, *SD*= 5.25). Data from four participants were unsuited for analysis: recordings from three participants were heavily artifact-laden, likely due to high impedance of the electrodes; data from one participant were corrupted by a failure to store event triggers, probably due to a memory issue with the smartphone. The final dataset for analysis comprised eight volunteers (6 female, 2 male; 1 left-handed, 6 right-handed, 1 ambidextrous) with an age range from 24 to 39 (*M* = 29.00, *SD*= 6.19). All participants provided written informed consent prior to their participation and were treated in accordance to the study protocol approved by the ethics committee of the University of Oldenburg.

### Paradigm

Participants performed auditory oddball tasks in the lab and in their office while following their normal office routines, including working on a computer, conversing with colleges, and going for lunch. The whole experiment took an average of 6.11 h (*SD* = 0.21 h) excluding the time to attach and remove the equipment (∼ 15 min).

Participants were equipped with a mobile ear-EEG setup for stimulus presentation and data acquisition (see apparatus for details and Figure 1A). The setup allowed them to move freely and to perform all of their normal office activities.

**Figure 1.**
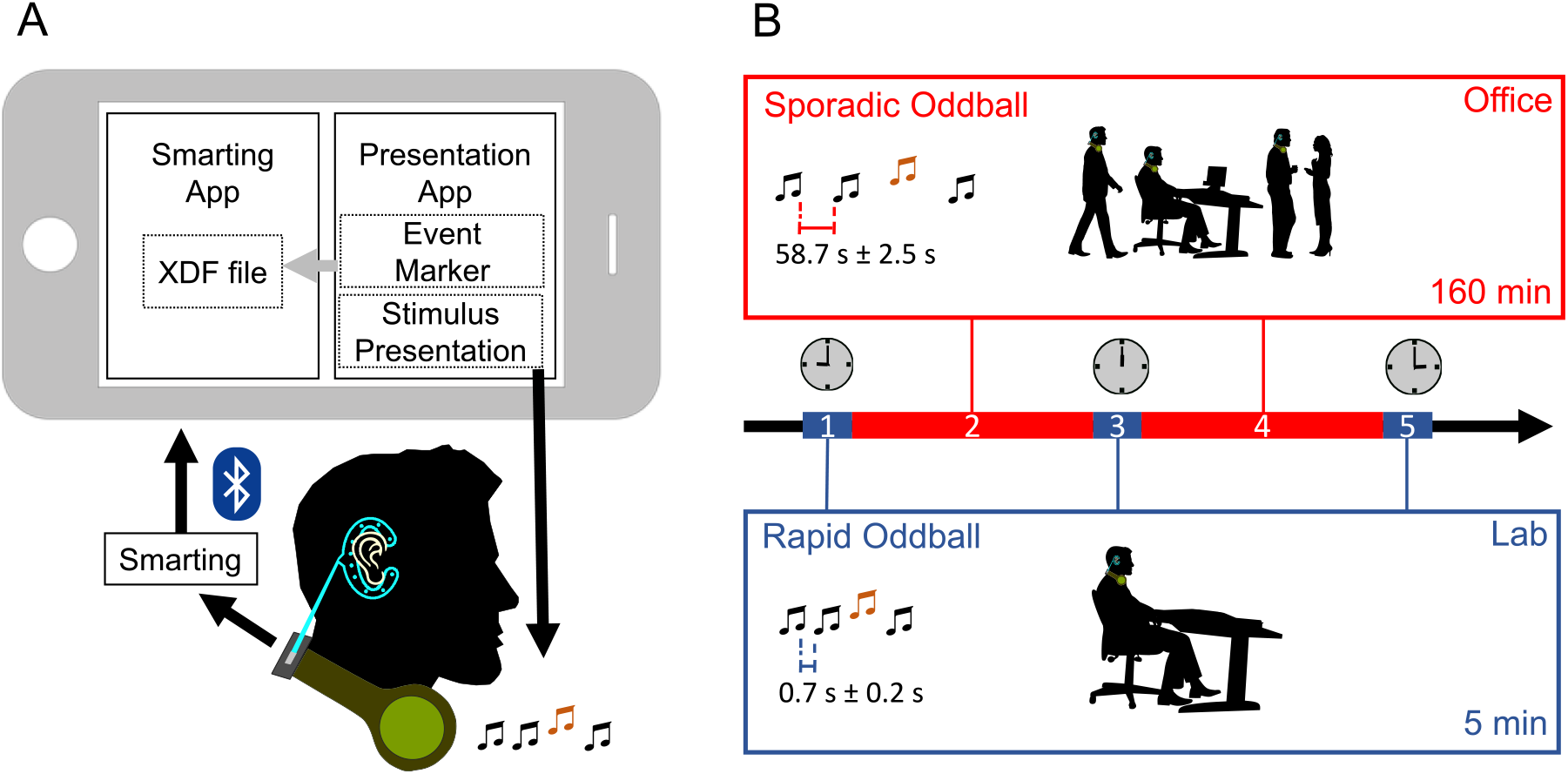
Illustration of experimental setup and procedure. **(A)** EEG was recorded with the cEEGrid. The amplifier was attached to headphones worn around the neck, which were also used to present the tones. A smartphone controlled stimulus presentation and stored the data. For stimulus presentation the Presentation app was used. The SMARTING amplifier transmitted to the recording smartphone via Bluetooth. The event markers and the amplifier data were synchronized by the SMARTING app running on the smartphone. **(B)** Participants completed 5 blocks: Blocks 1, 3, and 5 were rapid oddballs (5 min each) with interstimulus intervals of 500-900 ms. Participants performed this task while they were sitting quietly in a chair with no external disturbance either in the lab (block 1 and 5) or in their office (block 3). Block 2 and 4 were sporadic oddballs (160 min each) with sounds once approximately every minute. In this task participants were free to go about their normal office work. The recording was done continuously between ∼ 9AM and ∼ 15PM.

The auditory oddball task (Polich, 2007) was performed 5 times and came in two variants. The *rapid* variant lasted approximately 5 minutes and was administered three times (in the morning, around noon, and in the afternoon) with full focus on the task, either in the lab or in the participants’ office. In between these controlled recordings, the *sporadic* variant was administered for twice 160 minutes during regular office work. The sporadic variant differed from the rapid version by longer inter-trial intervals. The complete time course of the experiment is illustrated in Figure 1B.

In both versions, participants were presented with a sequence of double tones. Double tones were implemented to explore adaptation effects (cf. May & Tiitinen, 2010), but these will not be addressed in this paper. The first tone had a duration of 700 ms (including 5 ms rise and fall time) and was followed after a silent period of 100 ms by the second tone with identical pitch but a duration of 500 ms (including 5 ms rise and fall time). Two different double tones were used: an oboe-based sound with a fundamental frequency of 100 Hz and a clarinet-based sound with a fundamental frequency of 300 Hz. Tones were generated with Matlab (Version: 9.5.0.1049112; The Mathworks Inc., Natick, MA, USA) and had a sampling rate of 44100 Hz. Sound intensity was root mean square (RMS) normalized to achieve comparable intensity for all tones. One of the double tones served as target, the other as standard.

Target/standard assignment was counterbalanced across participants but constant across all 5 runs within each participant. The first four trials were always standards. The order of the remaining trials was randomized with no more than two targets in succession. Each run consisted of 160 trials (double tones), 112 standards (70%) and 48 targets (30%). Only targets required a response by the participant.

In the rapid oddball variant, the inter-trial interval (offset of previous double tone to onset of next double tone) randomly varied between 500 ms and 900 ms (*minimal standard random number generator* as implemented in Presentation), resulting in an average of 9 targets per minute. In the sporadic oddball, the inter-trial interval randomly varied between 56200 ms and 61200 ms (roughly once per minute) for an average of one target per 3.33 minutes. The infrequent sounds can be tolerated while concentrating on something else, making the task compatible with office work.

Participants were instructed to respond to target tones as fast as possible by touching the display of a smartphone which was attached to the upper arm of their non-dominant hand.

The sporadic oddball included 10 trials with short inter-trial intervals (alternating between targets and standards) at the beginning and at the end for familiarization and quality control. These trials were excluded from analysis. Moreover, after 80 sporadic trials in the sporadic condition were completed, the task was interrupted and participants were asked to sit still and rest for one minute with their eyes closed. This rest session served as a data quality check (see Figure 3 in the supplementary material).

### Apparatus

The setup consisted of headphones worn around the neck, a mobile EEG amplifier affixed to the headphones, two cEEGrids connected to the amplifier, a smartphone for stimulus presentation and data acquisition, and a power bank.

Consumer headphones (Sennheiser HD 4.50BTNC) served as speakers worn around the neck. They did not cover the ears. This allowed the participants to both hear the task sounds as well as all ongoing sounds around them. The tones were presented at maximum volume to be heard at a convenient loudness.

The experiment was programmed with Presentation (Version: 20.3; Neurobehavioral Systems, Inc., Berkeley, CA, USA) and stimulus presentation was performed by the Presentation Android app (Version: 2.0.9) running on a Sony Xperia Z3 Compact (OS: Android 6.0.1). The Presentation app also recorded hit rates and reaction times whenever the participants touched the smartphone display. The smartphone was rooted, all hardware-buttons (e.g., volume buttons) were disabled and a smart window cover (Sony Smartphone Cover SCR26) was used to block the software-buttons (e.g., home button). These smartphone modifications prevented accidental changes of the experimental settings by participants. To ensure sufficient battery life for the duration of the experiment (*>* 6.5 h), a power bank (Xoro MPB 255, 2500 mAh) was connected to the smartphone.

Two cEEGrids with ten Ag/AgCl electrodes apiece (tMSI, Oldenzaal, The Netherlands) were placed around the ears of the participants, one on each side. Electrode R4a served as driven-right-leg and R4b served as reference. The cEEGrids were connected to a mobile SMARTING 24-channel sleep EEG amplifier (mBrainTrain, Belgrade, Serbia) by a customized connector (see https://ceegrid.com). Impedance values of most electrodes were below 10 kΩ at the start of the experiment or approached this threshold during the course of the day. The amplifier, equipped with a 3D gyroscope and battery charge for at least seven hours, recorded with a resolution of 24 bits and a sampling rate of 250 Hz. Data were transmitted from the amplifier to the smartphone via Bluetooth and were recorded with the SMARTING Android app (Version: 1.7.0). The SMARTING app synchronized EEG data and the triggers from the oddball task using a Lab Streaming Layer framework (https://github.com/sccn/labstreaminglayer), producing an .xdf-file.

### Procedure

Participants arrived at the lab in the morning. The skin around their left and right ear was cleaned using abrasive gel and alcohol swabs. Small drops of electrolyte gel (Abralyt HiCl, EasycapGmbH, Germany) were applied to each electrode and the cEEGrids were positioned around the ear with double-sided adhesive tape. Participants were then equipped with the the rest of the setup. The amplifier was taped to a clip on the headphones, which were placed on participants’ neck. The cEEGrids were connected to the amplifier, the headphones and the power bank were connected to the smartphone. Smartphone and power bank were placed in a smartphone arm-pouch attached to the participants’ arm opposite their dominant hand. By attaching the smartphone to the participants’ arm, the display of the phone was conveniently accessible for participants to touch with the dominant hand (even during computer work). To ensure that the audio cable connecting the headphones and the smartphone was not accidentally disconnected, it was taped to the participants’ clothes on the back. The complete setup is shown in the supplementary material in Figure 1.

The experiment consisted of five blocks spanning just over six hours: a rapid block in the morning followed by a sporadic block, another rapid block at noon followed by the second sporadic block, and a last rapid block in the afternoon (see Figure 1B). Each block started with a one-minute calibration phase, where participants were asked to sit still and look at a fixation cross (see below). In each block, participants were first presented with their standard and then with their target tone. Participants initiated the start of the tone presentation by touching the display of the smartphone. The end of each block was announced by a speaker saying in German: “Done! Touch the display to complete the experiment.” Before the start of each block, the arm-pouch was removed to access the phone, store the recording to avoid data loss, and select the oddball variant (rapid/sporadic). The pouch was then reattached.

For the first block, participants were seated in front of a printed fixation cross that was placed on the wall approximately 80 to 100 cm away on eye level. The fixation cross served to reduce head movement, as participants had the tendency to orient their head towards the smartphone on their arm when pressing the display. Participants received written instructions and practiced the task with a short demo consisting of three standard tones and one target tone, which was followed by the one-minute calibration phase. The experimenter then left the room and participants completed the rapid oddball. Subsequently, participants received a new sheet with instructions for the sporadic oddball. After a short demo with one standard and one target and the one-minute calibration, the sporadic oddball started. During the 10 rapid trials in the beginning of the sporadic oddball, participants remained seated in the lab and after the first sporadic trial, they relocated to their office space, taking the written instructions with them. The instructions outlined the steps for the rest session, indicated which tone - high or low pitched - was the target, and provided contact information of the experimenter. After half of the sporadic oddball, the smartphone notified participants with the ringing of a service bell that it was time for the rest session. To stop the ringing, participants touched the display of the smartphone. To initiate the rest session, they touched the display again after closing their eyes. The rest session ended with another service bell notification, and with another display touch participants continued the main experiment.

At the end of the first sporadic oddball the experimenter met with the participants in their office for the second rapid oddball. Again, participants were provided with an instruction sheet and a fixation cross. The fixation cross, at eye level at an approximate distance of 80 to 100 cm, was either printed out and taped to a convenient object or it was presented on participants’ computer screen. Participants then completed the rapid oddball in their office, while the experimenter waited outside. In some cases, a (quiet) office mate was in the same room during this block.

Afterwards, the experimenter started the second sporadic oddball and left. For the last block - another rapid oddball - the experimenter and the participants met again in the lab. As before, participants received written instructions, were seated in front of the fixation cross, and completed the final block. Then the setup was removed and participants were provided with paper tissue to remove residual electrolyte gel.

During the sporadic oddball, participants were free to continue their everyday routines. They were instructed to respond to targets except when it would be unsafe to do so (e.g., when holding a cup with a hot beverage), and not to change or damage the experimental setup. Apart from office work on the computer, which was their predominant activity, this included drinking coffee, snacking, chatting with colleagues, going to lunch, or going for a small walk. Thus, the surrounding noise level varied over the course of the experiment. Some environments, such as the university cafeteria, were loud enough that the presented tones could not always be heard. However, this problem affected only a small portion of the recordings.

### Data Analysis

Data were analysed offline with Matlab (Version: 9.6.0.1335978; The Mathworks Inc., Natick, MA, USA) using and EEGLAB (Version: v2019.0; Delorme & Makeig, 2004) and custom scripts. All filters were zero-phase Hamming windowed sync finite impulse response filters as implemented in EEGLAB.

### Pre-Processing

A comprehensive pre-processing pipeline was implemented to reduce contamination of the in-lab and beyond-the-lab EEG by artifacts. The oddball data were low-pass filtered at 10 Hz (filter order 330) and high-pass filtered at 0.1 Hz (filter order 8250). The data were cleaned using Artifact Subspace Reconstruction as implemented in the EEGLAB plugin *clean_rawdata* (Version: 1.0; parameters: flatline criterion = 60, high-pass = [0.25 0.75], channel criterion = off, line noise criterion = off, burst criterion = 20, window criterion = off). ASR is a statistical anomaly detection method that compares statistical properties of clean calibration data with the properties of new data by using a series of linear subspace projections. With this procedure, artifacts, which typically induce high variance, are automatically identified and removed (cf. Mullen et al., 2015 for a detailed description of the ASR approach).

ASR is computationally efficient and can handle the complex artifacts occurring in mobile recordings (Blum et al., 2019; Dehais, Duprès, et al., 2019), making it especially well-suited for long-term mobile recordings. ASR does not rely on visual inspection, making it easier to reproduce than, for example, Independent Component Analysis.

The flatline rejection criterion of the ASR function rejected channels that contained a flat line for 60 s or longer. Remaining bad channels were identified and rejected by applying a liberal amplitude criterion (± 500 *µ*V) and then automatic channel rejection (*pop_rejchan*) based on the spectral properties of the channel, with a threshold of two standard deviations. This resulted in the complete rejection of one rapid block from one participant and one rapid and one sporadic block from a second participant. In these blocks, either all (1 sporadic) or at least half of all channels (1 rapid, 1 sporadic) were rejected. In the remaining blocks, on average 2.11 (*SD*= 2.75) channels were rejected across all participants. One rapid oddball block did not contain any event triggers and was also excluded. Rejected channels in the remaining datasets were then spherically interpolated. Altogether, 21 rapid oddball blocks (88%) and 15 sporadic oddball blocks (94%) remained for the final analyses.

Epochs were extracted from -0.2 to 0.8 s relative to the onset of the first tone in a double tone and baseline-corrected (−200 to 0 ms). The first two epochs in every block and missed targets (Rapid: *M* = 3.14, *SD*= 6.71; Sporadic: *M* = 5.67, *SD*= 5.07) were excluded. Bad epochs were identified and rejected using joint probability criteria as implemented in EEGLAB with a (global and local) threshold of two standard deviations. On average, 31% (*SD*= 7%) in the rapid oddball (standard: 69%; target: 31%) and 30% (*SD*= 4%) in the sporadic oddball (standard: 71%; target: 29%) were rejected. Time windows for extraction of mean P3 amplitude were defined as the latency of the peak amplitude ±100 ms in the range from 200 ms to 600ms after the first tone in a double tone in the per-condition grand average ERP. Mean P3 amplitude for each trial was extracted from 212 ms to 412 ms for the rapid oddball (peak latency 312 ms) and from 284 ms to 484 ms for the sporadic oddball (peak latency 384 ms).

### Gyroscope

Participants’ movement was recorded by the gyroscope in the amplifier. The amplifier was attached to headphones worn on the neck, therefore the gyroscope only measured gross body movement, in particular movements involving the shoulder area (i.e., head movements without involvement of the shoulder area were not necessarily captured). Displacement was calculated for each participant and block by taking the square root of the sum of the squared gyroscope axes 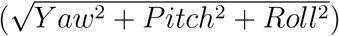. This value is the absolute angular acceleration independent of direction, which can be interpreted as the total deviation from the initial position.

Individual displacement thresholds were calculated by combining data from all three rapid blocks and taking the median displacement plus four times the interquartile range.

### Statistical Analysis

Statistical analysis was performed with RStudio (Version: 1.2.5042; RStudio, Inc., Boston, MA, USA; R-Version: 3.6.3). For all tests, an alpha level of 0.05 was used (Bonferroni-corrected in case of multiple comparisons).

### Behavioural Data

Behavioural data were analysed using generalized linear mixed models (GLMM) with the R-packages *lme4* (Bates, Mächler, Bolker, & Walker, 2015) and *lmerTest* (Kuznetsova, Brockhoff, & Christensen, 2017). GLMMs were calculated to take inter-individual differences into account and to test the hypotheses regarding between-condition differences in reaction times and response accuracy (hit/miss). The GLMM for reaction times included the fixed factors *oddball variant* (rapid/sporadic), a random intercept for *participant*, and a by-participant random slope for *oddball variant*. This model was fit with several distributions and link functions to find the combination best suited for the data. We report results from the best fit (an inverse Gaussian distribution with an inverse link) as assessed by likelihood ratio tests (Baayen, Davidson, & Bates, 2008; Baayen & Milin, 2010; see supplementary Tables 1-3 for full results including likelihood ratio tests). The GLMM for response accuracy similarly included *oddball variant* as fixed factor, a random intercept for *participant*, and a by-participant random slope for *oddball variant*. This model was fit with a binomial distribution and a logit link function. The statistical significance of differences in reaction times or hit rates between oddball variants under the ‘null hypothesis’ of no difference was evaluated with the Wald chi-square test.

### EEG Data

For statistical analysis of ERPs we selected the mean of channels R2 and R3 re-referenced to the mean of channels R6 and R7 (henceforth: vertical bipolar cEEGrid channel) based on previous work with the cEEGrid (Bleichner & Debener, 2017; Bleichner, Mirkovic, & Debener, 2016; Debener et al., 2015), suggesting this signal may be expected to yield the highest amplitudes. Due to fluctuations in the number of trials per condition and participant, we used linear mixed models (LMM) to assess differences in P3 amplitudes between standard and target tones for each condition. We used a LMM with independent random intercepts and random slopes (dependence between intercepts and slopes made the model non-identifiable). The statistical significance of differences standard and target tone amplitudes under the ‘null hypothesis’ of no difference was evaluated with the Wald chi-square test.

### Gyroscope Data

Displacement per block was compared using one-way repeated measures ANOVA followed by pairwise post-hoc t-tests where appropriate. We report generalized eta squared 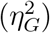 as a measure of effect size.

## Results

We analyzed behavioural data, EEG data, and movement data. Altogether, we collected a total of 47.40 hours of data, of which 41.66 hours were suitable for analysis (ranging from 3.11 hours to 6.35 hours per participant). This total time comprises 2.66 h from the rapid oddball and 39.01 h from the sporadic oddball.

### Behavioural Data

Behavioural data were available from all five blocks for each of the 8 participants included in the EEG analysis. The GLMM with inverse Gaussian distribution and inverse link function (see supplementary Table 1) yielded an estimated mean reaction time of 1.09 seconds (*β*_*link*_ = 0.91, *SE*_*link*_ = 0.06) for the rapid oddball. The mean increase in reaction time from the rapid to the sporadic oddball was 0.83 seconds (*β*_*link*_ = -0.39, *SE*_*link*_ = 0.05). Participants responded significantly faster (*χ*^2^(1) = 65.31, *p <* .001) in the rapid than in the sporadic oddball. Results from GLMMs using other distributions and link functions yielded substantively similar results (see supplementary Table 2; results of other GLMMs not shown). The observed reaction times per block are illustrated in Figure 2A.

**Figure 2.**
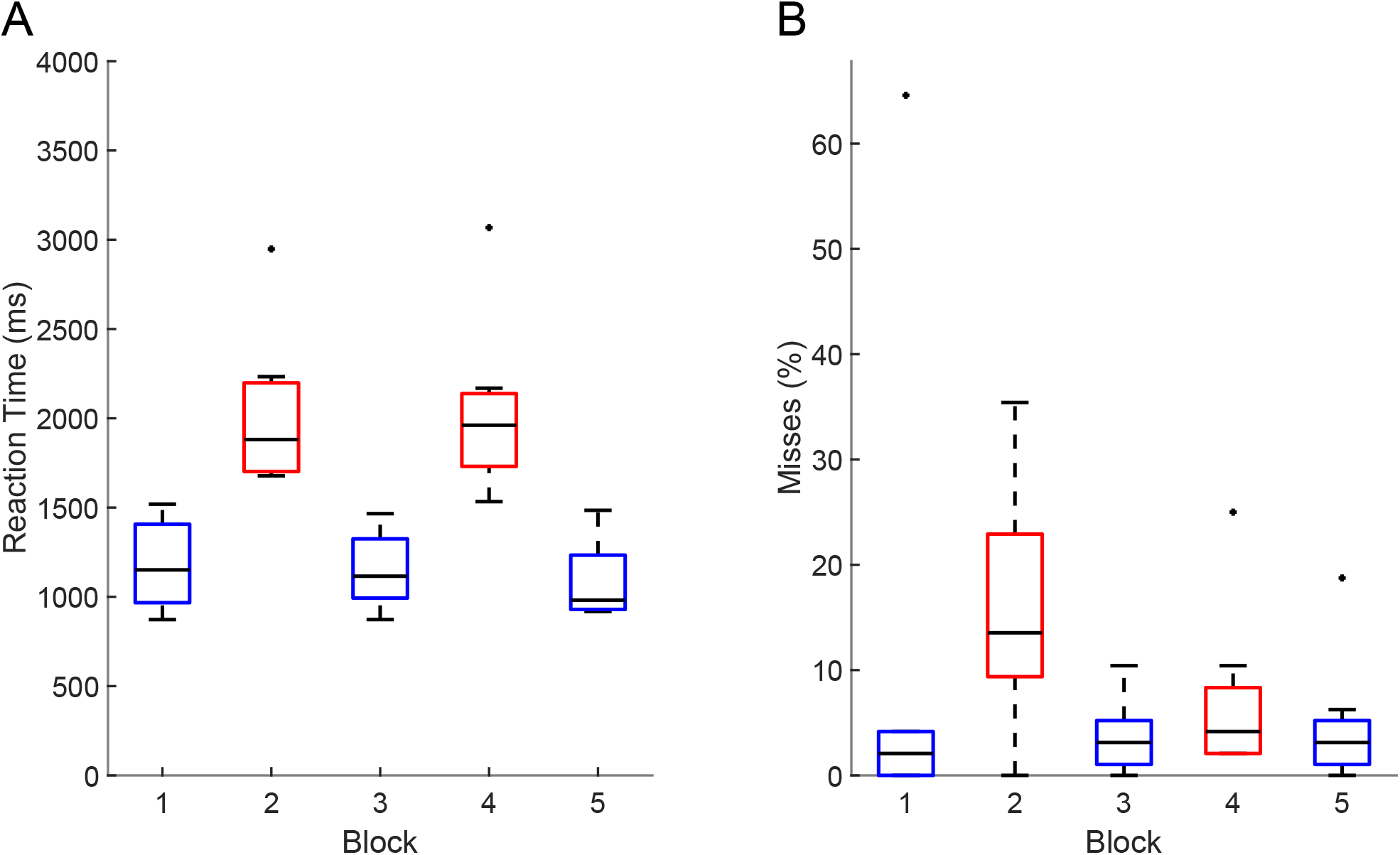
**(A)** Average reaction times per block. **(B)** Average proportion of misses per block. Blue: rapid oddball; red: sporadic oddball. The central black line indicates the median, the bottom and top edges indicate the 25^th^ and 75^th^ percentile. Whiskers indicate the most extreme data points not considered outliers which are indicated by a black dot. Outliers were defined as data points deviating more than 1.5 times the interquartile range from the bottom or top edges.

**Figure 3.**
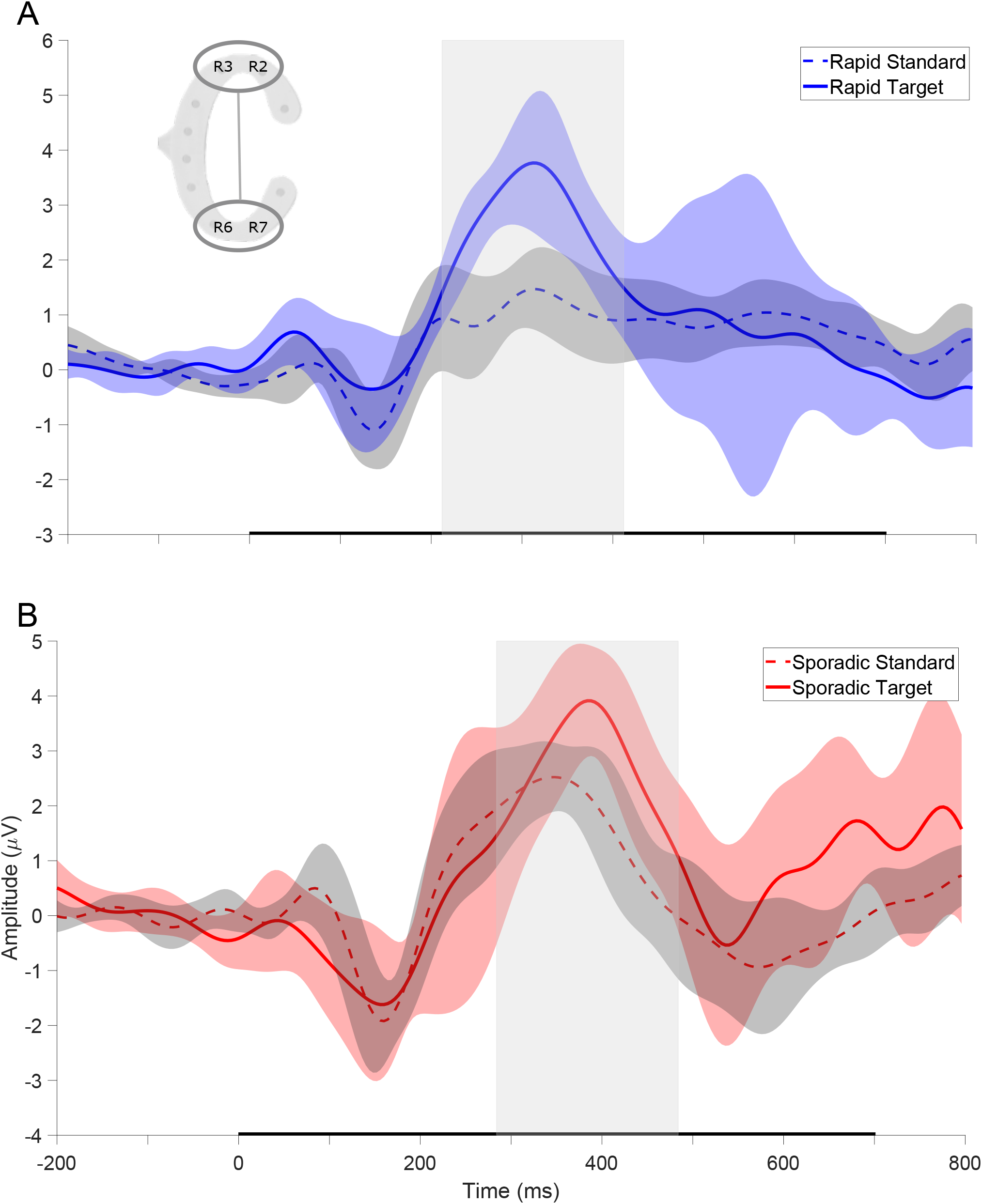
Grand average ERPs of the vertical bipolar channel (channel pair illustrated in the top left corner, (R3+R2)/2 - (R6+R7)/2). **(A)** Rapid oddball. **(B)** Sporadic oddball. Dotted lines indicate standard tones, continuous lines indicate target tones. The black bar on the x-axis represents tone presentation. The central grey box shows the time window used to extract the P3 amplitude. The shaded areas indicate the 95% confidence interval.

The mean chance of missing a target in the rapid oddball was 3.5% (*β*_*link*_ = -3.31, *SE*_*link*_ = 0.43) and increased by 6.9% (*β*_*link*_ = 1.15, *SE*_*link*_ = 0.39) in the sporadic oddball (see supplementary Table 3). The chance of missing a target was significantly higher in the sporadic than in the rapid oddball (*χ*^2^(1) = 8.94, *p <* .01). This difference is mainly driven by the sporadic block where most participants went to lunch and could thus hardly hear the tones (see procedure). Figure 2B shows the observed percentage of misses per block.

### EEG Data

Figure 3 shows the grand average ERP for both oddball variants for the vertical bipolar channel. The ERP for both conditions is characterized by an early negative component between 100 and 200 msec and a later positive component between 200 and 500 msec (P3). Please refer to supplementary Figure 2 for single channel grand average ERPs and supplementary Tables 4 and 5 for detailed results of the LMM analyses. In the rapid oddball, the mean amplitude for the standard tone was 1.07 *µ*V (*SE* = 0.22) with mean increase of 1.75 *µ*V (*SE* = 0.31) for the target tone. The Wald chi-squared test indicated that this difference was significant (*χ*^2^(1) = 30.92, *p <* .001). In the sporadic oddball, the mean amplitude for the standard tone was 1.63 *µ*V (*SE* = 0.27) with a mean increase of 1.20 *µ*V (*SE* = 0.33) for the target tone. This difference was also significant ((*χ*^2^(1) = 13.211, *p <* .001). In sum, both conditions showed significantly higher ERP amplitudes in response to target than to standard tones (see also Figure 4).

**Figure 4.**
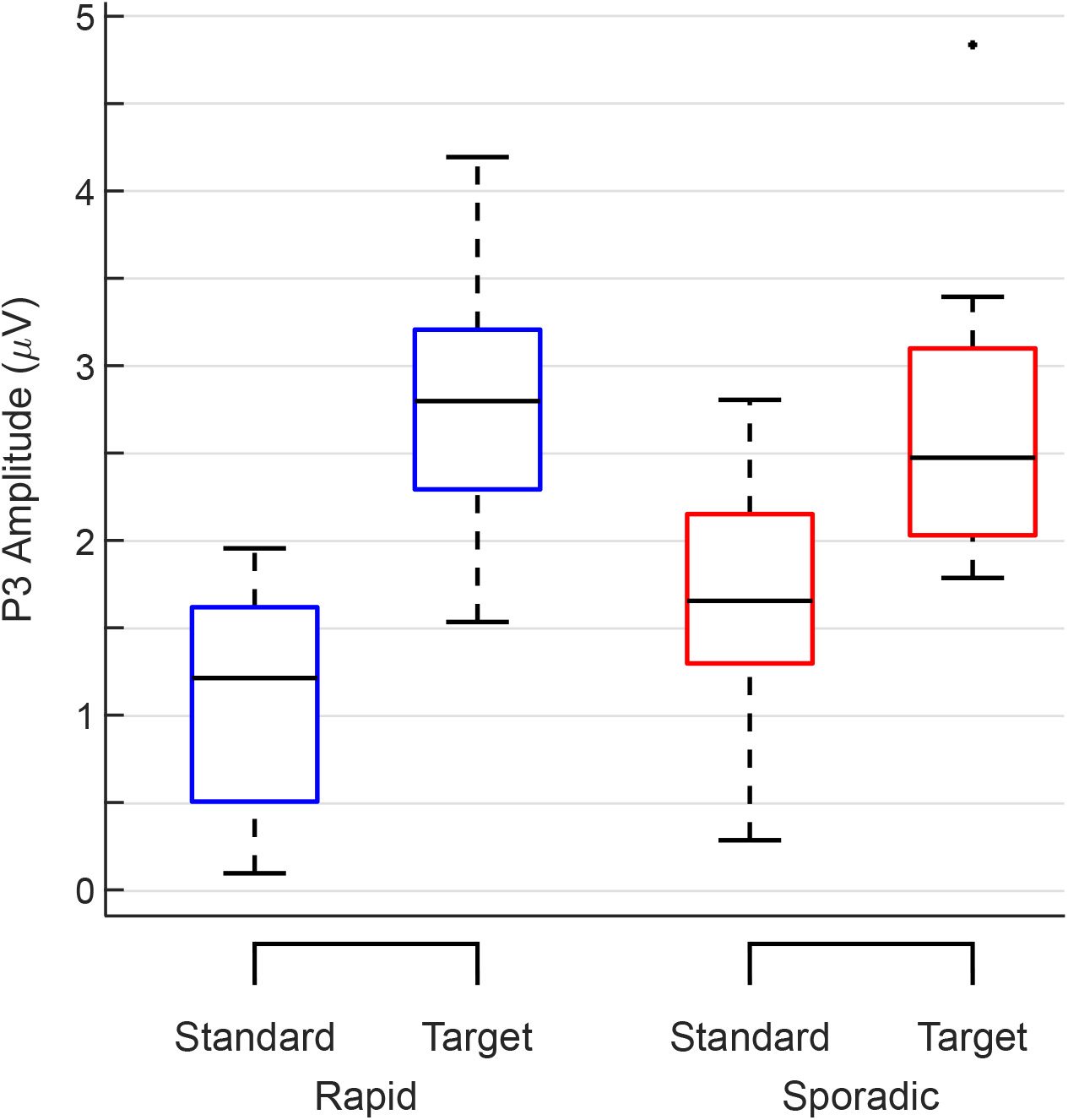
Boxplots of ERP amplitudes. P3 amplitudes per type of tone and condition. Blue: rapid oddball; red: sporadic oddball. The central black line indicates the median, the bottom and top edges indicate the 25^th^ and 75^th^ percentile. Whiskers indicate the most extreme data points not considered outliers, which, in turn, are indicated by a black dot. Outliers were defined as data points deviating more than 1.5 times the interquartile range from the bottom or top edges.

### Gyroscope Data

Figure 5A displays overall displacement per block, averaged across all participants. For each block and participant, the percentiles of the displacement value were calculated and then averaged. As expected, more movement (i.e., higher displacement values) occurred in the sporadic oddball blocks (Block 2 and Block 4) than in the rapid oddball blocks (Block 1,3, and 5). Nevertheless, comparison with the global threshold (dashed magenta line) demonstrates that even in the sporadic blocks (solid and dashed red lines) most of the data - approximately 75% - were free of gross movement, compared to about 95% in the rapid blocks (for which the participants were instructed to sit down). The boxplot inset shows mean displacement per block (please refer to supplementary Figure 4 for axis-specific movement). A repeated measures ANOVA for displacement with the factor *Block* yielded a significant main effect (*F* (1,4) = 34.01, *p <* .001, *η*^2^ = .80). Post-hoc t-tests revealed that all rapid blocks (1,3,5) differed significantly (all *p <* .001) from all sporadic blocks (2,4). Differences between rapid blocks did not reach significance, whereas the difference between the two sporadic blocks was significant (*p* = .04).

**Figure 5.**
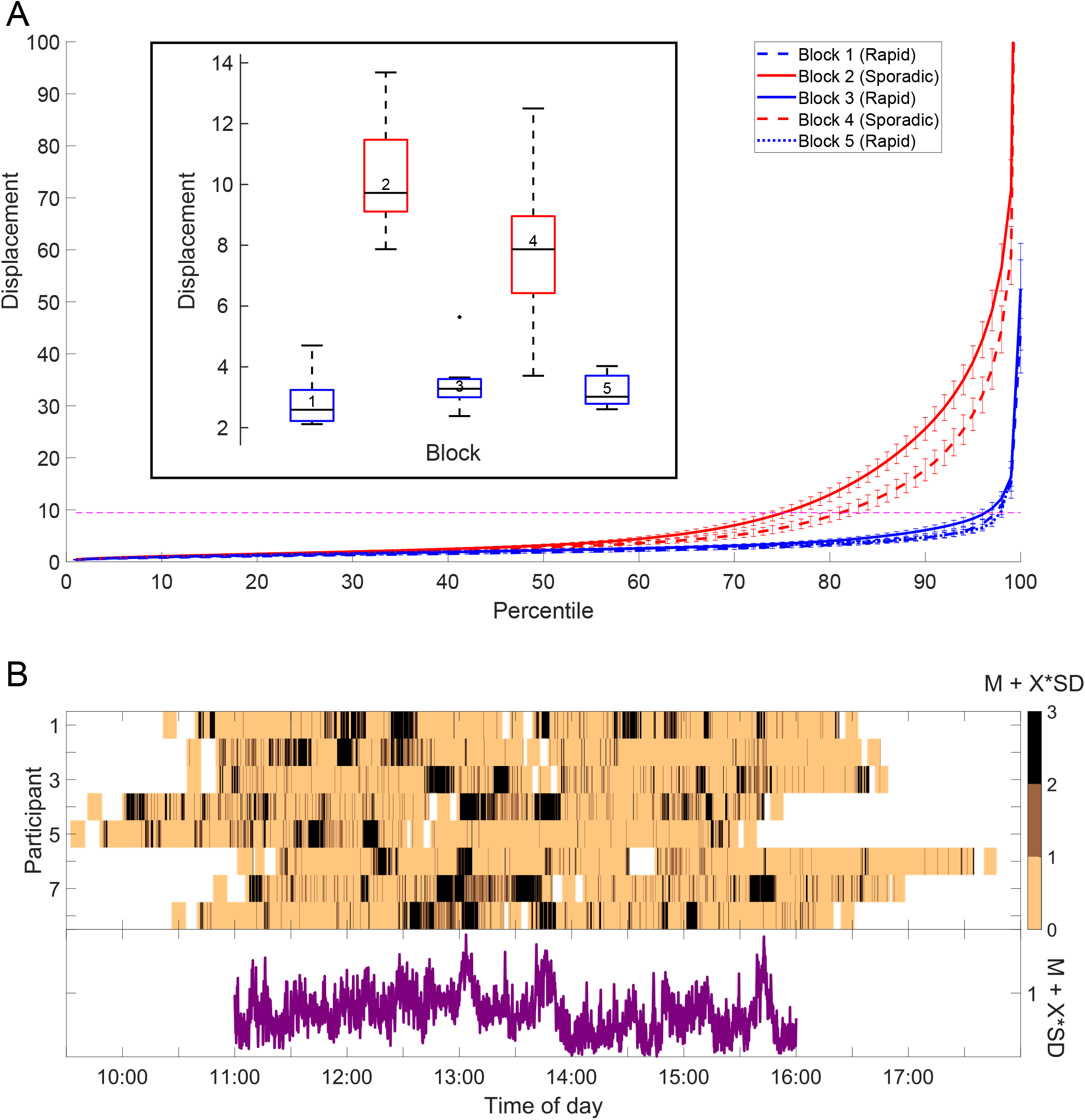
**(A)** Distribution of displacement per block. Shown is the displacement at each percentile. Higher displacement values indicate more movement. Error bars represent 95% confidence intervals. The magenta line indicates the global threshold (mean of all individual thresholds) of what was considered as relative rest. The boxplot inset shows displacement per block. Blue: rapid oddball; red: sporadic oddball. The central black line indicates the median, the bottom and top edges indicate the 25^th^ and 75^th^ percentile. Whiskers indicate the most extreme data points not considered outliers, which, in turn, are indicated by a black dot. Outliers were defined as data points deviating more than 1.5 times the interquartile range from the bottom or top edges. **(B)** Time course of movement data. The upper panel shows the amount of movement over time for each participant.The movement has been categorized for each participant based on their displacement mean and standard deviation from all three rapid blocks. The brightest color indicates movement within one standard deviation plus the participant-specific mean. The darker colors represent increments of the standard deviation added to the mean. This plot has been smoothed with a moving median over 10 seconds. The lower panel shows the grand average of the movement categorization. The average is only shown between 11:00 AM and 16:00 PM where data from most participants (∼7) were available at the same time. Due to different start and end times, the recordings do not perfectly overlap.

In the upper panel, Figure 5B shows the time course of movement data categorized based on participant-specific mean and standard deviation of the displacement. The lower panel shows the corresponding averaged time course. The peaks around 13:00 PM and 13:45 PM indicate when most participants left for and returned from lunch (i.e., walking to and from the cafeteria).

## Discussion

We adapted a lab-based paradigm for real-world situations to investigate selective auditory attention in an everyday context using mobile ear-EEG. The auditory oddball ran in the background while participants worked in a normal office environment. The aim of this work was to demonstrate the feasibility of EEG long-term recordings in everyday scenarios to study auditory attention and auditory perception. Behaviourally, participants were able to discriminate between target and non-target sounds while going about their regular office routines. Similarly, the observed differences in P3 amplitude imply differences in brain responses to target and standard stimuli during office work.

With rare exceptions, hardware and software proved durable and reliably measured brain activity for six hours. Gyroscope data allowed us to trace the participants’ activity on a coarse level. Despite periods of movement, participants spent most of their day at relative rest (∼ 75%), suggesting limited influence of large movement on EEG signal quality. All participants reported that the sporadic oddball did not interfere notably with their office activities. Few misses show that participants performed the sporadic oddball adequately, even while focusing on their work. All participants reported that the setup was comfortable to wear throughout the day, though they felt relieved when the equipment was removed at the end of the day.

Reactions times and misses were higher in the sporadic than in the rapid oddball. These results mirror dual-task studies, where participants are instructed to perform a primary and a secondary task, such as watching a movie while performing an oddball task (e.g., Willard, Johnson, & Rosenfeld, 1994). In the sporadic oddball participants similarly followed their work routine while performing the oddball task. Participants are slower and make more mistakes in dual-task conditions than when performing only a single task (e.g., Karatekin, Couperus, & Marcus, 2004). Our results therefore confirm our hypothesis and suggest that performing an experimental task beyond the lab may generally be conceived of as dual-task situations where participants dynamically regulate their allocation of resources between the experimental task and their self-chosen other task. In future studies, an independent measure of how strongly the participants are engaged in either task may help to assess trade-off effects.

Unlike classical lab experiments, beyond-the-lab studies in everyday life have to deal with numerous uncontrolled factors, for example, the natural soundscape, social encounters, and the individuals’ choice of the current activity. These factors influence auditory perception in everyday situations. The loss of experimental control is therefore inevitable and makes each recording unique. Here we analysed reaction times and response accuracy using GLMMs to capture variance at the inter- and intra-participant level and to test hypotheses. However, our model ignored uncontrolled factors. In future studies, gaining a fuller understanding of auditory perception in everyday situations requires that we characterize these influences and include them in the analysis. Ultimately, a multi-modal approach will be necessary to gather information about the participants’ current state and their surroundings using a multitude of sensors, as implemented in lab-based mobile applications (Gramann et al., 2014). Consequently, multivariate statistics beyond those we employed here are required to interpret these datasets comprehensively.

The ERP analysis demonstrated higher P3 amplitudes for target than for standard tones (Polich, 2007) in both the rapid and the sporadic oddball, replicating previous work using ear-EEG (Debener et al., 2015; Denk et al., 2018). Here we have extended the results of Debener and colleagues (2015) who had participants wear cEEGrids for a whole day, but only recorded data in the mornings and in the afternoons. In between, the cEEGrid was affixed to participants’ ear but not connected to an amplifier. In the present study, in contrast, brain activity was recorded continuously during a whole day. Hence, the present study showed that cEEGrids, coupled with reliable equipment, cannot only be worn but also measure brain activity for a whole day.

Recordings beyond the lab come with technical and methodological challenges. Experimental paradigms designed for the lab cannot necessarily be transplanted beyond the lab, especially when we are interested in “natural behaviour". Instead, we need to find a compromise between naturalness and sufficient experimental control. In the present study, the rapid oddball was adapted in such a way that it could be performed while working. The time interval between tones was substantially increased to reduce interference with office activities. This change did indeed make it feasible to perform this task while doing office work, but it also diminished the habituation to the standard tone which occurs when the tones are presented in quick succession. In a conventional rapid oddball, this habituation makes detection of deviant target tones automatic and effortless (Polich, 2007). In the sporadic oddball the detection of the target tones may not have been automatic due to the long time intervals between tones. Both standard and target tones were isolated events that stood out against the current soundscape.

Each tone had to be compared to some internal copy of the target sound (memory trace). In other words, standard tones potentially demanded more cognitive evaluation in the sporadic than in the rapid oddball. This difference may be reflected in the seemingly higher amplitudes for the standard tone in the sporadic task compared to the rapid one. This points to the important caveat that repeated presentation of a stimulus in classical lab studies may alter our perception and may not generalise to the varied soundscapes we encounter in daily life.

The present paradigm artificially imposed the relevance of tones by assigning standard and target tones. Analysis of brain responses to auditory stimuli that are ecologically meaningful to participants, such as their personal ringtones or their names (Perrin et al., 2005; Roye, Schröger, Jacobsen, & Gruber, 2010) may increase ecological validity even further. Such stimuli could serve as targets whereas other ringtones or names of strangers could serve as standards. The P3 response to one’s own name could then be contrasted with other names. By recording over several days using transparent EEG (Bleichner & Debener, 2017) it may even be possible to rely entirely on responses to naturally occurring events.

Acquisition of behavioral data could be improved as well. Participants had longer reaction times to targets in the sporadic than in the rapid oddball, since during the sporadic oddball they were engaged in other tasks as well. This difference is informative insofar as it reflects the time participants needed to stop their current activity (e.g., typing on the keyboard or using the mouse) and press the display of the smartphone. To measure reaction times with less interference in their current activity, a double eye blink or a voice command could be used in the future.

### Conclusion

We recorded ear-EEG data for six hours during an office day. Our results demonstrated that it is feasible to study auditory attention using mobile ear-EEG, highlighting the potential of beyond-the-lab experimentation. We discuss the technical, procedural and methodological pitfalls of adapting classical laboratory paradigms to a real-world context. This study helps to pave the way for a fuller understanding of auditory perception in everyday contexts. Our work adds to the growing number of studies that show the general feasibility of beyond-the-lab EEG recordings (e.g., Debener et al., 2012; Ladouce, Donaldson, Dudchenko, & Ietswaart, 2019; Scanlon et al., 2019; Wascher et al., 2016), but more work is required to use the richness of this methodology.

## Acknowledgement

We thank Björn Holtze and Stefan Debener for their feedback and ideas. We thank Reiner Emkes for his technical support. This work was funded by the Deutsche Forschungsgemeinschaft (DFG, German Research Foundation) under the Emmy-Noether program - BL 1591/1-1 - Project ID 411333557.

## Open practices statement

The datasets are available from the corresponding author on request. The experiment was not preregistered.

## Supplementary material

**Supplementary Figure 1.**
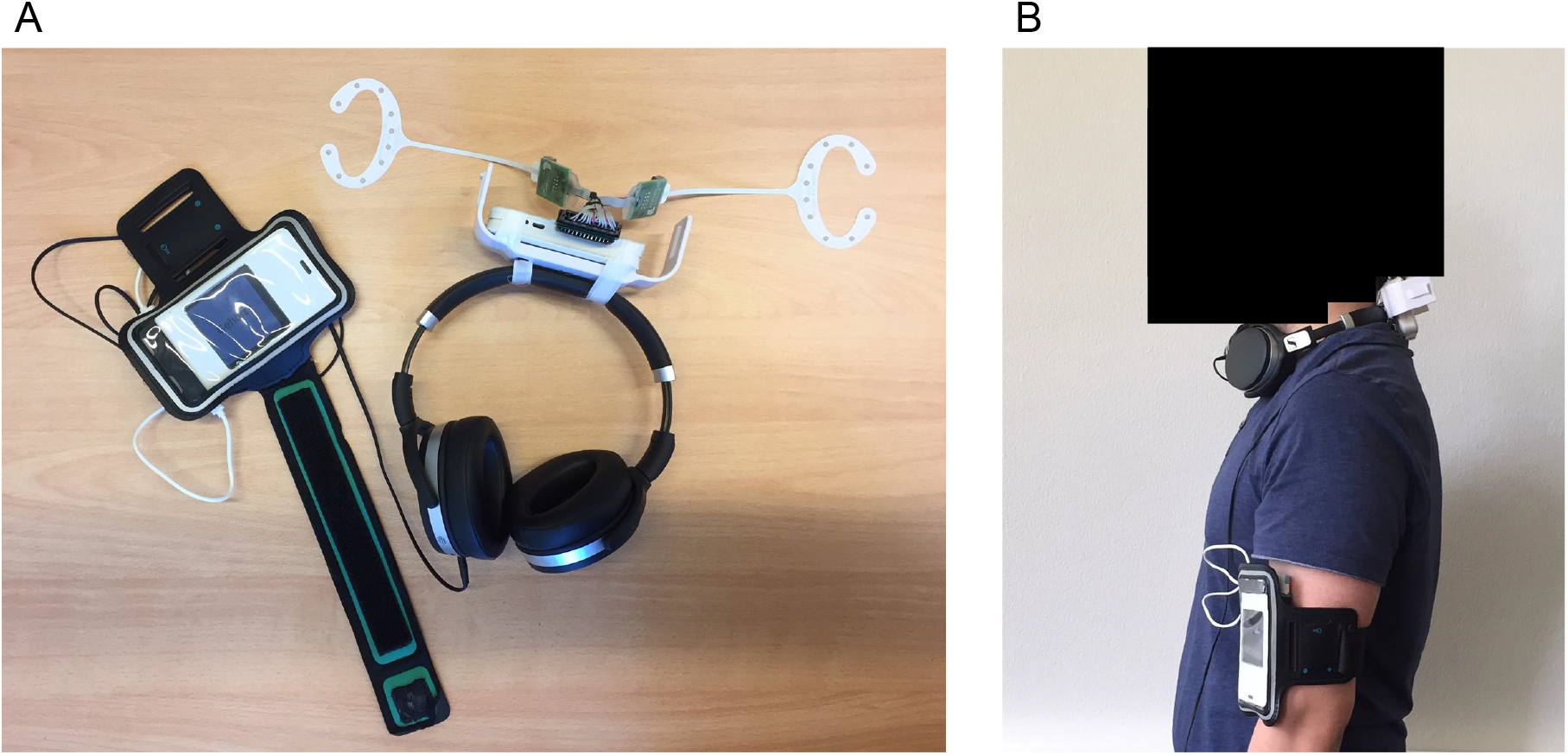
Complete experimental setup. **(A)** cEEGrids were connected to the amplifier, the amplifier was taped to the headphones, and the headphones were connected to the smartphone via an audio cable. A power bank (not visible) was connected to the smartphone and both were stored in an arm-pouch. The smartphone both recorded data and presented the sounds. **(B)** Participant wearing the setup. The headphones were worn around the neck and the smartphone was attached to the arm.

**Supplementary Figure 2.**
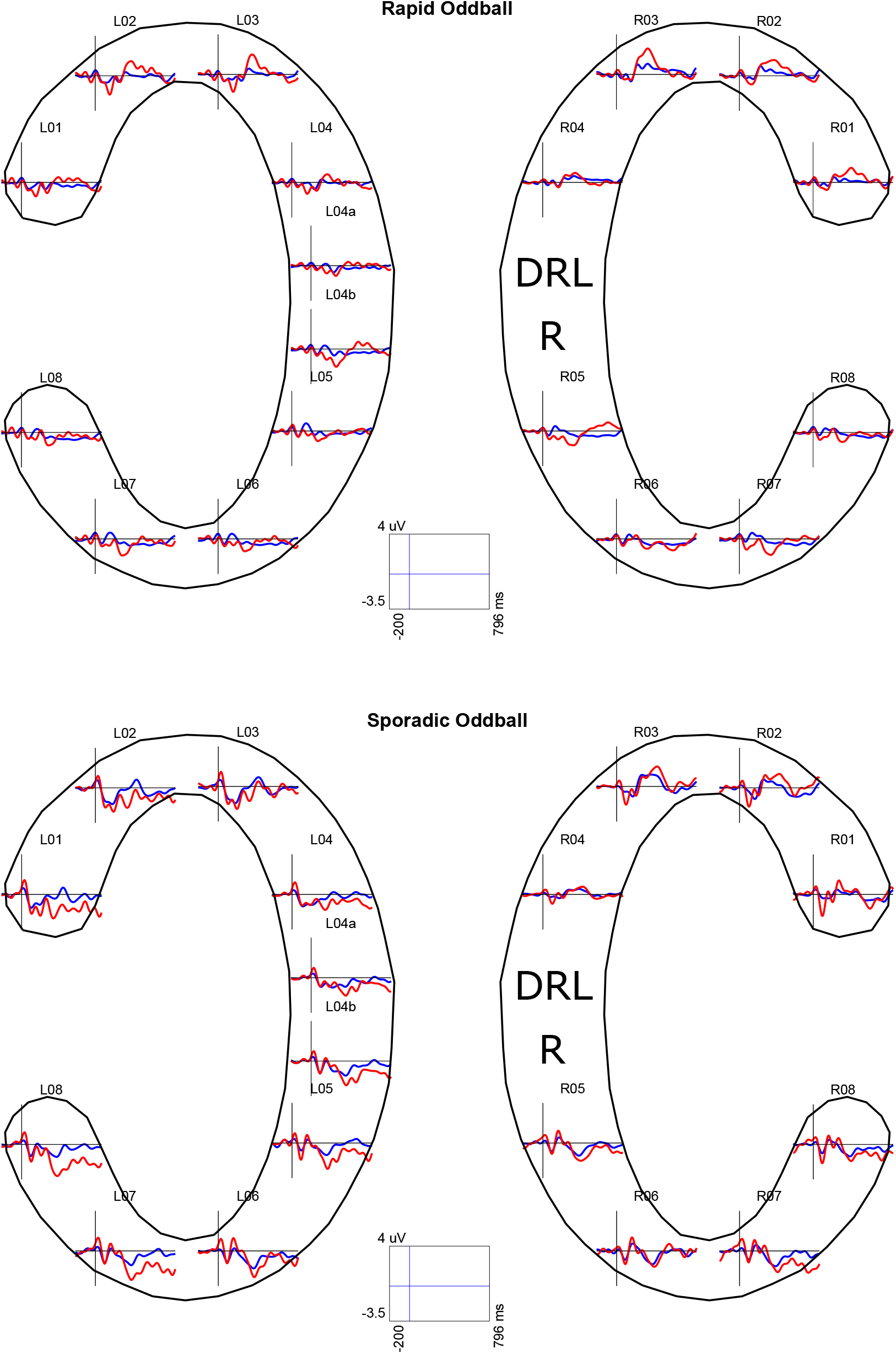
Grand average of channel ERPs shown in the cEEGrid layout including electrode labels. Standard tones are blue and target tones are red. DRL: Driven-right-leg; R: Reference electrode. Top: Grand average ERPs from the rapid oddball. Bottom: Grand averaged ERPs from the sporadic oddball.

**Supplementary Figure 3.**
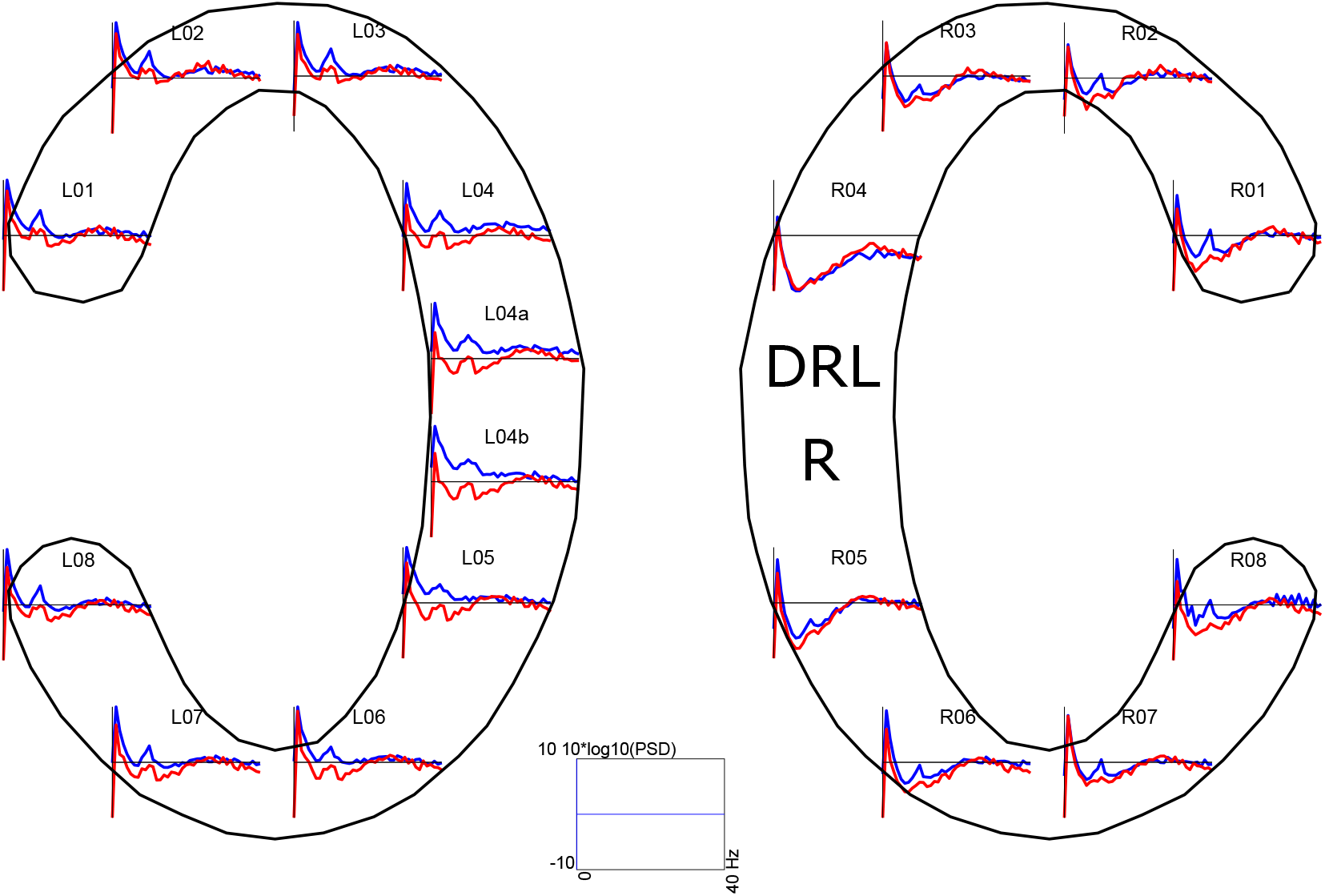
Grand average of EEG spectra for the rest session in the sporadic oddball shown in the cEEGrid layout including electrode labels. Blue: eyes closed; red: eyes open; DRL: Driven-right-leg, R: reference electrode. *Note on data processing*: The data from the rest session of the sporadic oddball was low-pass filtered at 40 Hz (filter order 166) and high-pass filtered at 1 Hz (filter order 1650). The same pre-processing pipeline as the ERP analysis was applied (except the filters). For this analysis, the data from the rest session served as eyes closed condition and the data from the calibration phase in the beginning of a block served as eyes open condition. Both conditions had a duration of one minute. From each of those conditions, consecutive windows comprising 1024 samples with an overlap of 60 samples were extracted and Hanning windowed. These windows were submitted to a fast Fourier transform (Pwelch as implemented in Matlab) and subsequently averaged and log-normalized (10*log10). Note that both conditions originated from different time points in the experiment that are 80 minutes apart, reducing their comparability; however, the rest session only served as a data sanity check and was not statistically evaluated.

**Supplementary Figure 4.**
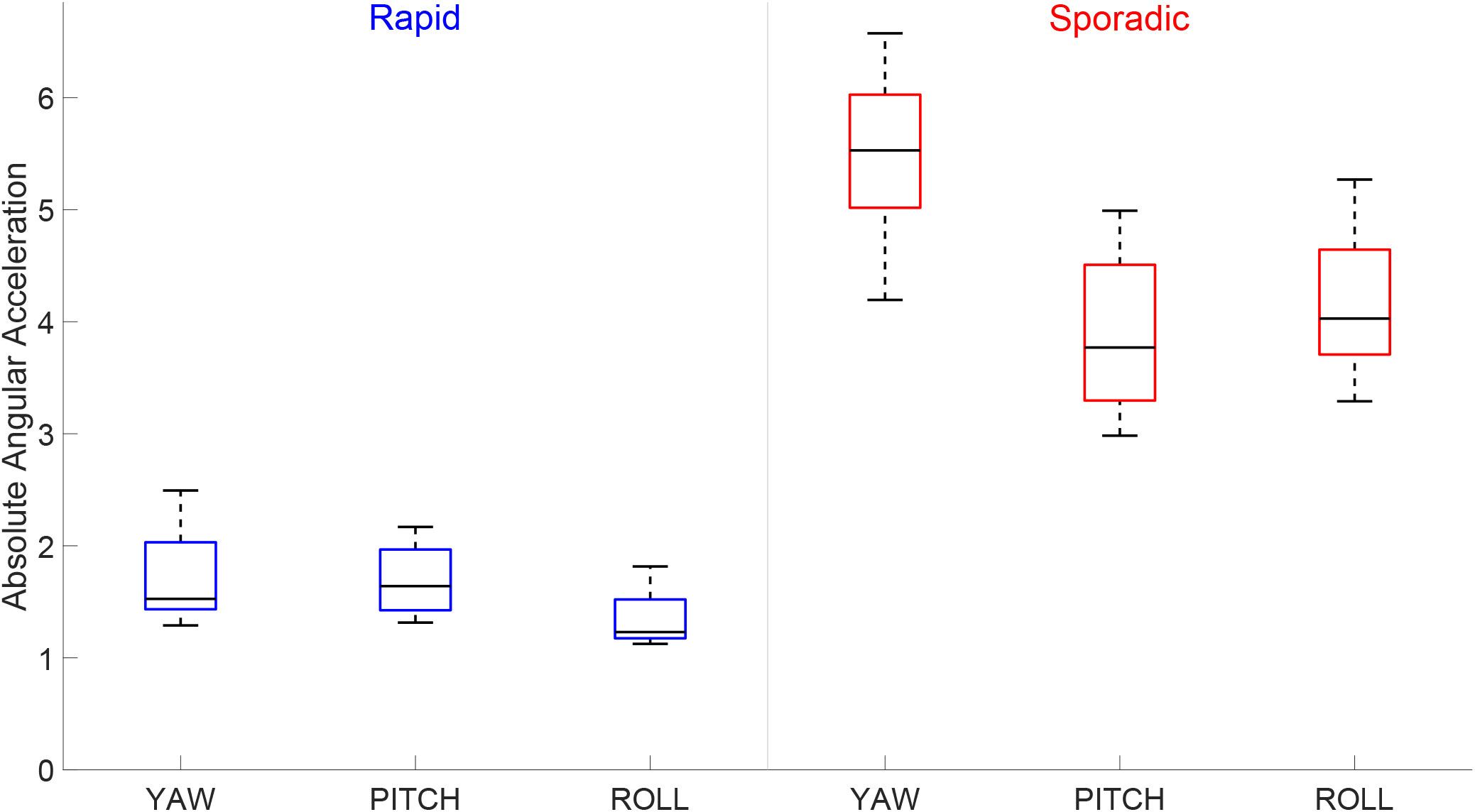
Absolute values of angular acceleration per condition and gyrospcope axis. The central black line indicates the median, the bottom and top edges indicate the 25^th^ and 75^th^ percentile. Whiskers indicate the most extreme data points not considered outliers, which, in turn, are indicated by a black dot. Outliers were defined as data points deviating more than 1.5 times the interquartile range from the bottom or top edges. *Note on statistics*: These data were submitted to a two-way repeated measures ANOVA with the factors *condition* (rapid/sporadic) and *axis* (yaw, pitch, roll). A significant main effect of condition, *F* (1,7) = 125.95, *p <* .001, *η*^2^ = .88, and axis, *F* (2,14) = 38.76, *p <* .001, *η*^2^ = .35, was found. The interaction was also significant, *F* (2,14) = 49.47, *p <* .001, 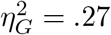. Post-hoc tests revealed that in the sporadic condition, both pitch and roll significantly differed from yaw (*p <* .01). Furthermore, the corresponding axes from both conditions also differed significantly from each other (all *p <* .001).

**Supplementary Table 1.** *GLMMs for reaction times*

| Fixed Effects | Model Summary |  |  |  |
| --- | --- | --- | --- | --- |
| | $\beta$ | SE | t-value | p-value |
| Intercept | 0.91 | 0.06 | 14.40 | <.001 |
| Oddball Variant | -0.39 | 0.05 | -8.08 | <.001 |

| Random Effects |  |  |  |
| --- | --- | --- | --- |
|  | Name | Standard Deviation | Correlation |
| Participant | Intercept | 0.07 |  |
|  | Slope | 0.07 | -0.80 |

*Note:* GLMM was fitted with an inverse Gaussian distribution and an inverse link. Coefficients, standard errors, and standard deviations are not back-transformed. The intercept is the predicted mean reaction time in *seconds*<sup>-1</sup> for the rapid oddball; the beta for oddball variant represents the predicted mean change (on the link scale!) when moving from the rapid to the sporadic oddball.

**Supplementary Table 2.** *Model comparison for reaction times*

| | AIC | BIC | log-likelihood | $\chi^2$ | df | p |
| --- | --- | --- | --- | --- | --- | --- |
| F: Gaussian, L: Identity | 4088.9 | 4121.8 | -2038.5 |  |  |  |
| F: Gaussian, L: Log | 4081.5 | 4114.4 | -2034.8 | 0 | 0 | ns |
| F: Gaussian, L: Inverse | 4080.1 | 4112.9 | -2034.0 | 0 | 0 | ns |
| F: Gamma, L: Identity | 1290.4 | 1323.2 | -639.2 | 0 | 0 | ns |
| F: Gamma, L: Log | 1287.0 | 1319.8 | -637.5 | 0 | 0 | ns |
| F: Gamma, L: Inverse | 1285.1 | 1317.9 | -636.5 | 0 | 0 | ns |
| F: Inverse Gaussian, L: Identity | 700.9 | 733.7 | -344.43 | 3388.1 | 0 | <.001 |
| F: Inverse Gaussian, L: Log | 697.3 | 730.2 | -342.7 | 3384.2 | 0 | <.001 |
| F: Inverse Gaussian, L: Inverse | 695.1 | 728.0 | -341.6 | 3385.0 | 0 | <.001 |

*Note:* F = Family; L = Link function; ns = non-significant

**Supplementary Table 3.** *GLMM for response accuracy*

| Fixed Effects | Model Summary |  |  |  |
| --- | --- | --- | --- | --- |
| | $\beta$ | SE | z-value | p-value |
| Intercept | -3.31 | 0.43 | -7.61 | <.001 |
| Oddball Variant | 1.15 | 0.39 | 2.99 | <.01 |

| Random Effects |  |  |  |
| --- | --- | --- | --- |
|  | Name | Standard Deviation | Correlation |
| Participant | Intercept | 1.10 |  |
|  | Slope | 0.88 | -0.88 |

*Note:* GLMM was fitted with a binomial distribution and a logit link function. Coefficients, standard errors, and standard deviations are not back-transformed and represent the predicted log odds of *missing* a target. The intercept is the predicted mean log odds of missing for the rapid oddball; the beta for oddball variant is the predicted mean change in log odds when moving from the rapid to the sporadic oddball.

**Supplementary Table 4.** *LMM for P3 amplitudes rapid oddball*

| Fixed Effects | Model Summary |  |  |  |
| --- | --- | --- | --- | --- |
| | $\beta$ | SE | t-value | p-value |
| Intercept | 1.07 | 0.22 | 4.84 | <.01 |
| Type of tone | 1.75 | 0.31 | 5.56 | <.001 |

| Random Effects |  |  |
| --- | --- | --- |
|  | Name | Standard Deviation |
| Participant | Intercept | 0.51 |
|  | Slope | 0.59 |

*Note:* The intercept is the predicted mean ERP amplitude ( $\mu\text{V}$ ) for the standard tone. The beta for type of tone is the predicted mean change in ERP amplitude when moving from the standard tone to the target tone (i.e., the difference in amplitude between the tones).

**Supplementary Table 5.** *LMM for P3 amplitudes sporadic oddball*

| Fixed Effects | Model Summary |  |  |  |
| --- | --- | --- | --- | --- |
| | $\beta$ | SE | t-value | p-value |
| Intercept | 1.63 | 0.27 | 5.93 | <.001 |
| Type of tone | 1.20 | 0.33 | 3.64 | <.01 |

| Random Effects |  |  |
| --- | --- | --- |
|  | Name | Standard Deviation |
| Participant | Intercept | 0.62 |
|  | Slope | 0.32 |

*Note:* The intercept is the predicted mean ERP amplitude ( $\mu\text{V}$ ) for the standard tone. The beta for type of tone is the predicted mean change in ERP amplitude when moving from the standard tone to the target tone (i.e., the difference in amplitude between the tones).

## Notes

### Competing Interest Statement

The authors have declared no competing interest.

### Summary of Updates

Confidence intervals in figure 3 were corrected

